# Characterization of *N*-glycosylation and its Functional Role in SIDT1-Mediated RNA Uptake

**DOI:** 10.1101/2023.08.01.551569

**Authors:** Tingting Yang, Haonan Xiao, Xiulan Chen, Le Zheng, Hangtian Guo, Jiaqi Wang, Xiaohong Jiang, Chen-Yu Zhang, Fuquan Yang, Xiaoyun Ji

## Abstract

The mammalian SID-1 transmembrane family members, SIDT1 and SIDT2, are multi-pass transmembrane proteins that mediate the cellular uptake and intracellular trafficking of nucleic acids, playing important roles in the immune response and tumorigenesis. Previous work has suggested that human SIDT1 and SIDT2 are *N*-glycosylated, but the precise site-specific *N*-glycosylation information and its functional contribution remain unclear. In this study, we employ high-resolution liquid chromatography tandem mass spectrometry to comprehensively map the *N*-glycosites and quantify the *N*-glycosylation profiles of SIDT1 and SIDT2. Further molecular mechanistic probing elucidates the essential role of *N*-linked glycans in regulating cell surface expression, RNA binding, protein stability, and RNA uptake of SIDT1. Our results provide crucial information about the potential functional impact of *N*-glycosylation in the regulation of SIDT1-mediated RNA uptake and provide insights into the molecular mechanisms of this promising nucleic acid delivery system, with potential implications for therapeutic applications.

## INTRODUCTION

Mammalian SID-1 transmembrane family member 1 (SIDT1) and member 2 (SIDT2) are the human orthologs of *Caenorhabditis elegans* systemic RNA interference defective protein 1 (SID-1), which is required for transporting exogenous double-stranded RNA (dsRNA) into the cytoplasm and is therefore essential for systemic RNA interference in *Caenorhabditis elegans* (1-4). Previous studies on mammalian SIDT1 and SIDT2 have implicated them in various biological processes such as glucose and lipid metabolism, immune response, and tumorigenesis, including breast, lung, gastrointestinal, pancreatic and non-small cell lung cancers (NSCLC), but the specific mechanisms are unknown (5-13). In contrast, the role of SIDT1 and SIDT2 in regulating various physiological processes by mediating cellular RNA uptake and intracellular trafficking have been extensively studied, providing significant insights into their functional contributions.

SIDT1 mediates the internalization of extracellular microRNA (miRNA) and small interfering RNA (siRNA) and subsequent gene silencing, which has great therapeutic potential for a wide range of human diseases, such as cancer, viral infections, and cardiovascular diseases (14-18). Our previous studies showed that the acidic environment enhances RNA transport efficiency via SIDT1, and provided a potential molecular basis for SIDT1 mediated RNA uptake (18, 19). Similarly, SIDT2 mediates gymnosis, facilitating the uptake of naked single-stranded oligonucleotides into living cells (20). In addition to being involved in extracellular RNA uptake, both SIDT1 and SIDT2 exhibit lysosomal membrane localization and play a crucial role in the activation of the innate immune response by transporting internalized dsRNA across the lysosomal membrane to engage the cytoplasmic retinoic acid-inducible gene I (RIG-I)-like receptors (RLRs) and thereby induce type I interferon (IFN-I) production (10, 11, 21). SIDT2 also mediates the delivery of cellular DNA and RNA into lysosomes for degradation through an arginine-rich motif that binds to RNA/DNA (22, 23). However, the precise link between the physiological processes associated with SIDT1 and SIDT2 and their critical role in nucleic acid transport activities remains unclear.

Sequence- and structure-based analyses showed that most members of the mammalian SID-1 transmembrane family are multi-pass membrane proteins consisting of eleven transmembrane helices (TMs) and a large N-terminal extracellular domain (ECD) (19, 24). The ECD contributes to SIDT1- and SIDT2-dependent RNA transport and their direct binding to RNA (19, 25). Both SIDT1 and SIDT2 have a potential zinc-binding site with three histidine residues within the TMs responsible for lipolytic activities (19, 24, 26, 27). In addition to RNA binding, previous biochemical and structural data have suggested that human SIDT1 and SIDT2 are *N*-glycosylated (19, 24, 28, 29). Glycosylation is one of the most abundant and diverse forms of post-translational modification of proteins that is common to all eukaryotic cells (30). Numerous roles have been ascribed to protein *N*-glycosylation (31). In particular, *N*-glycans can intrinsically modulate protein folding and stability, provide protection against proteases, and mediate protein-ligand interactions. Proper protein *N*-glycosylation is also crucial for the formation of protein complexes, the modulation of protein-protein interactions and the correct assembly of higher order protein structures (32-35).

The mammalian SID-1 transmembrane family-dependent nucleic acid transport has been demonstrated in several *in vitro* and *in vivo* systems (10, 11, 15, 17, 18, 20, 22, 23). However, a clear understanding of the role of its *N*-glycosylation in this process has not been achieved. Further sequence analysis revealed that the *N*-glycosylation of SID1 transmembrane family members is conserved across mammals (Figure S1). The high conservation of *N*-glycosylation argues for an essential biological function, and it seems reasonable to assume that *N*-glycosylation plays an essential role in the mammalian SID-1 transmembrane family. We hypothesized that *N*-glycosylation of the mammalian SID-1 transmembrane family has important implications for RNA transport, RNA binding, protein stability, etc.

Here we report the first systematic characterization of site-specific *N*-glycosylation and the corresponding *N*-glycans of human SIDT1 and SIDT2 using high-resolution liquid chromatography-tandem mass spectrometry (LC-MS/MS). We demonstrate that human SIDT1 relies on its *N*-glycosylation to achieve its correct functional localization. We also found that deficient *N*-glycosylation affects the RNA binding, protein stability, and RNA uptake of SIDT1. Taken together, these observations unravel the *N*-glycosylation puzzle and reveal the importance of the conserved *N*-glycosylation of the mammalian SID-1 transmembrane family for RNA transport, RNA binding, proper functional localization, etc. These results may help to elucidate the regulatory pathways of the cellular function of the SID-1 transmembrane family.

## RESULTS

### The mammalian SID-1 transmembrane family is *N*-glycosylated

We searched for evolutionarily conserved (Asn-X-Ser/Thr, where X is any amino acid except for Pro) motifs in the amino-acid sequences of the mammalian SID-1 transmembrane family from different species using NetNGlyc 1.0 (36). The prediction results show that the mammalian SID-1 transmembrane family contains 7-10 potential *N*-glycosylation sites. Of these sites, 3-4 are located between the TMs, while the remaining sites are located in the ECD (based on human SIDT1 sequence, shown in Figure 1A). However, whether these sites are *N*-glycosylated remains unknown. To verify the *N*-glycosylation status of human SIDT1, we first heterologously overexpressed SIDT1 in human embryonic kidney (HEK293T) cells and treated the cell lysates with Peptide-*N*-Glycosidase F (PNGase F) to study the enzymatic deglycosylation effect. As shown in the immunoblotting result, SIDT1 was found to have an approximate mass of 100 kDa. As expected, after treatment with PNGase F, the migration of SIDT1 was accelerated and the apparent molecular mass decreased to approximately 90 kDa (Figure 1B). Similar results were obtained when we performed the same experiments with SIDT2, indicating that SIDT1 and SIDT2 are *N*-glycosylated (Figure 1B), in agreement with previous studies (28, 29).

**Figure 1.**
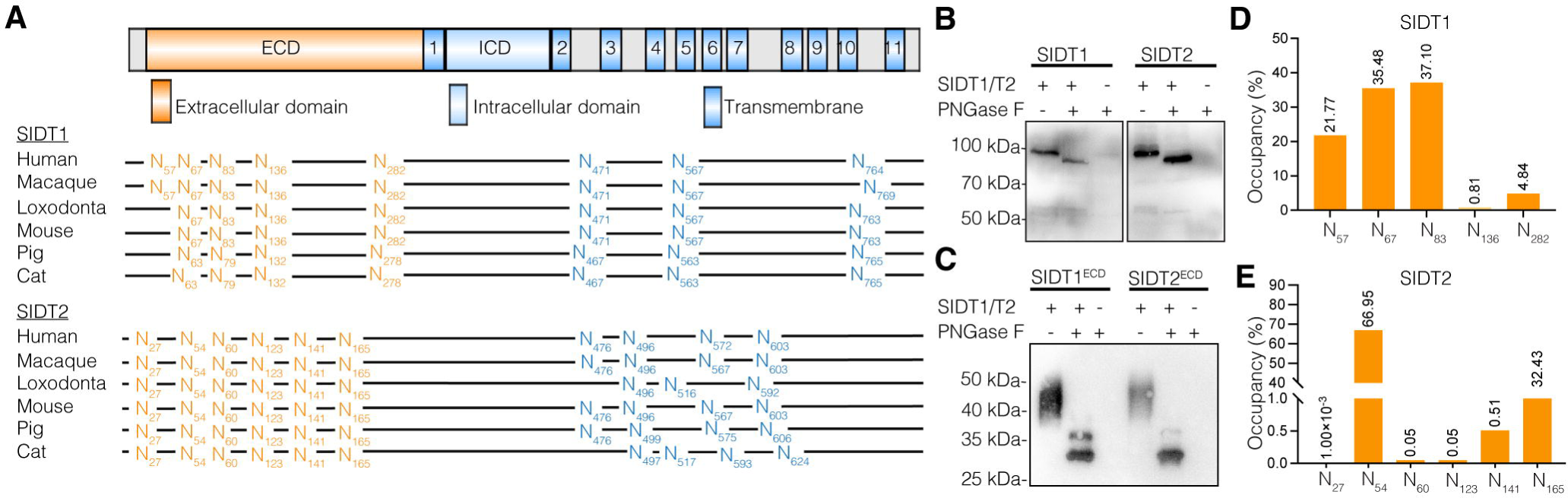
The mammalian SID-1 transmembrane family is *N*-glycosylated. **A** Domain schematic of human SIDT1 (upper panel) and conservation of *N*-glycosylation sites in mammalian SIDT1 and SIDT2 based on sequence alignment (bottom panel). **B** DeL*N*Lglycosylation analysis of full-length SIDT1 and SIDT2 with (+)/without (-) PNGase F, analyzed using Western blot with anti-Flag antibody. **C** DeL*N*Lglycosylation analysis of SIDT1^ECD^ and SIDT2^ECD^ with (+)/without (-) PNGase F, analyzed using Western blot with anti-His antibody. **D, E** Quantification of the abundance of glycopeptides at each glycosylation site of SIDT1 (**D**) and SIDT2 (**E**) through high-resolution LC-MS/MS analysis. Three independent experiments were performed. ECD, extracellular domain; PNGase F, peptide-N-glycosidase F.

Considering that *N*-glycosites of SIDT1 and SIDT2 are mainly located in the ECD, we also recombinantly obtained the truncated proteins (SIDT1^ECD^ and SIDT2^ECD^) from mammalian cells (HEK293F) and evaluated their *N*-glycosylation status. Immunoblot analysis reveals a broad band at approximately 50 kDa, and the width of the electrophoretic band is characteristic of glycosylated proteins and can be attributed to the heterogeneity of the carbohydrate moieties (Figure 1C). After treatment with PNGase F, SIDT1^ECD^ and SIDT2^ECD^ appeared as a single 30 kDa band, and the apparent molecular mass decreased by ∼20 kDa (Figure 1C).

To determine the site-specific *N*-glycosylation of SIDT1, the SIDT1 protein recombinantly expressed in HEK293F cells with intact *N*-glycosylation was digested into peptides and the intact glycopeptides were analyzed with LC-MS/MS using EThcD mode. The workflow for site-specific *N*-glycosylation characterization of recombinant proteins is shown in Figure S2. We identified five extracellularly localized glycosylation sites (N57, N67, N83, N136, N282) out of the eight predicted *N*-glycosites of SIDT1, and the remaining three intra-transmembrane-located sites were unoccupied. The occupancy of glycopeptides at each glycosylation site is shown in Figure 1D. The *N*-glycosylation modification occurred mainly at N57, N67 and N83, as these three sites accounting for over 90% of the identified glycopeptides. Consistent with SIDT1, no intra-transmembrane-located *N*-glycosites were detected in SIDT2. Six *N*-glycosites (N27, N54, N60, N123, N141, N165) were identified in the ECD of SIDT2 and the occupancy of each glycosite is shown in Figure 1E. Moreover, over 99% of the identified *N*-glycopeptides were mainly from N54, N141 and N165.

Taken together, these results suggest that SIDT1 and SIDT2 are highly *N* glycosylated, solely within the ECD.

### Site-specific *N*-glycosylation analysis of recombinant SIDT1 and SIDT2

Intact *N*-glycopeptide analysis can provide comprehensive glycoprotein information, including the composition and number of *N*-glycans decorating a specific *N*-glycopeptide or *N*-glycosite. Hundreds of nonredundant intact *N*-glycopeptides were identified from the recombinant SIDT1 and SIDT2 ECDs expressed in HEK293F cells (supplementary Tables. S1 and S2). The top twelve *N*-glycans appearing at different glycosites of SIDT1 and SIDT2 were mostly of the complex type terminating with sialic acid (NeuAc) or with fucosylation (Figure 2A). Glycoproteins are heterogeneous with a high degree of glycan microheterogeneity. SIDT1 and SIDT2 were decorated with 149 and 133 unique *N*-glycan compositions on the five and six *N*-glycosites, respectively (Figure 2B). Different *N*-glycosites have different degrees of glycan heterogeneity. On SIDT1^ECD^, three *N*-glycosites (N57, N67, and N83) have high glycan heterogeneity with a high proportion of complex/hybrid type glycans. 26, 102, and 94 glycan compositions were observed on the three *N*-glycosites, respectively. The other two *N*-glycosites (N136 and N282) have lower glycan heterogeneity, with only 9 and 13 glycan compositions on each glycosite, respectively. A similar degree of glycosylation was observed for SIDT2^ECD^. Three *N*-glycosites (N54, N141, and N165) have a high degree of glycan heterogeneity. 99, 42, and 78 glycan compositions were observed on the three glycosites, respectively, while other three *N*-glycosites (N27, N60, and N123) have limited glycan heterogeneity, with only 4, 8, and 3 glycan compositions on the three glycosites, respectively.

**Figure 2.**
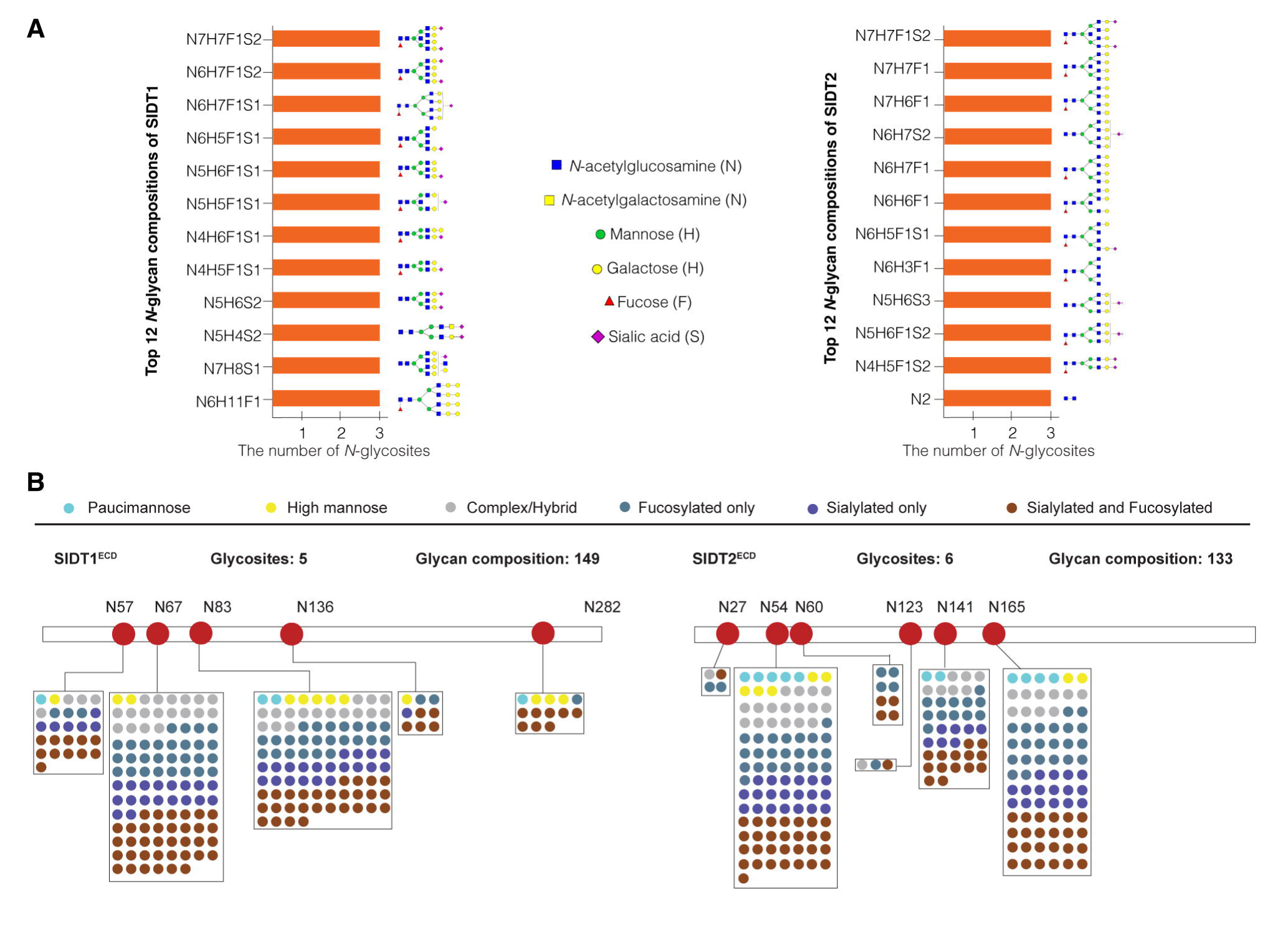
Site-specific *N*-glycan occupancy of recombinant SIDT1 and SIDT2. **A** The top twelve *N*-glycan compositions at the three major sites of the recombinant SIDT1 (left panel) or SIDT2 (right panel). **B** Different types and *N*-glycan compositions on each *N*-glycosite of the recombinant SIDT1 (left panel) or SIDT2 (right panel).

In all, our data reveal a regular *N*-glycan occupancy on mammalian SID-1 transmembrane family, with a remarkable heterogeneity in *N*-glycan compositions. This meta-heterogeneity of glycosylation in SIDT1 and SIDT2 might link to specific function (37).

### *N*-glycosylation is a prerequisite for the proper cell surface expression of SIDT1

To characterize the effects of the *N*-glycan, we generated SIDT1^ECD^ mutant protein (SIDT1^ECD^ mutant) with all Asn (N) to Gln (Q) substitutions of the *N*-glycosylation sites. The small-scale expression studies were performed at the shake flask level to elucidate the effect of *N*-glycosylation on SIDT1^ECD^ protein secretion. We found that wild-type SIDT1^ECD^ folds and assembles correctly in HEK293F cells and is robustly secreted into the media. In contrast, SIDT1^ECD^ mutant caused a significant and substantial secretion defect (Figure 3A). Notably, site-directed single amino acid mutation of each *N*-glycosylation site resulted in the N57Q, N67Q and N136Q SIDT1^ECD^ variants being secreted abnormally and retained intracellularly (Figure 3B and Figure S3). This suggests that *N*-glycosylation is critical for successful folding and secretion of SIDT1^ECD^, indicating that *N*-glycosylation is essential to ensure correct protein localization of SIDT1.

**Figure 3.**
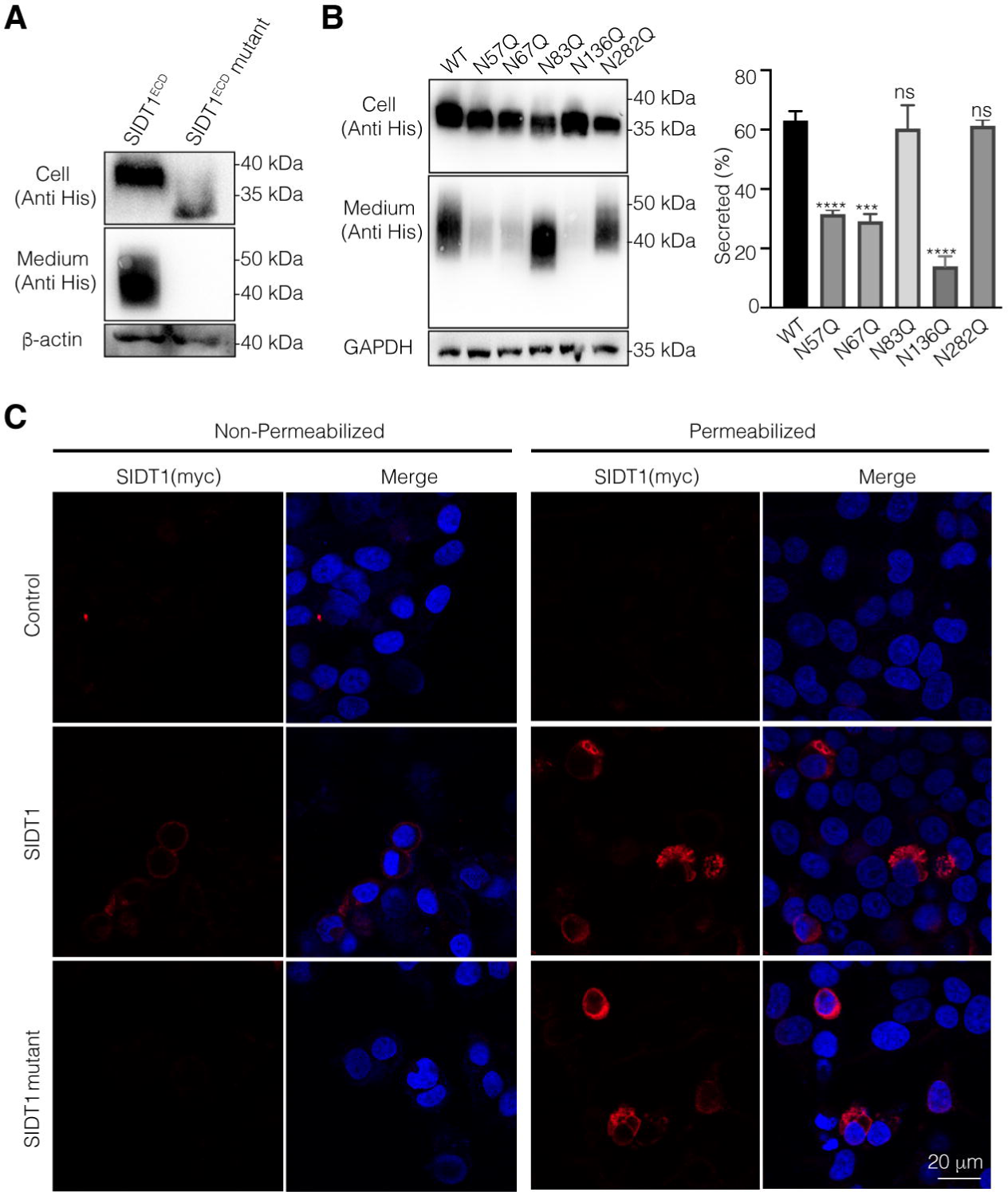
*N*-glycosylation is important for cell surface expression of SIDT1. **A** The effect of *N*-glycosylation on the secretion of wild-type SIDT1^ECD^ and SIDT1^ECD^ mutant (cell and medium), and protein levels were analyzed by Western blot. **B** Secretion of the wild-type (WT), N57Q, N67Q, N83Q, N136Q and N282Q SIDT1^ECD^ variants was analyzed by Western blot. Band density was quantified using ImageJ software (right panel). Data are represented as means ± S.D, and statistical significance is determined by One-way ANOVA. ns, not significant, ***P < 0.001, ****P < 0.0001. **C, D** Cell surface expression of wild-type SIDT1 and SIDT mutant variant was determined by indirect immunofluorescence in non-permeabilized (**C**) or permeabilized (**D**) HEK293T cells by probing with anti-myc antibody. DAPI (blue) was used for nuclear staining. Scale bar, 20 μm. Three independent experiments were performed. DAPI, 2-(4-amidinophenyl)-6-indolecarbamidine dihydrochloride; ECD, extracellular domain; HEK293T, human embryonic kidney.

Next, we tested the effect of *N*-glycosylation on the subcellular localization of SIDT1. We examined the wild-type SIDT1 and SIDT1 mutant for cell surface expression by confocal microscopy in the absence and presence of saponin (a detergent that permeabilizes cell membranes without disrupting them). The myc-tagged SIDT1 variants were transiently transfected into HEK293T cells to assess SIDT1 expression at the cell membrane using an antibody against myc as a marker for the membrane surface, since the antibody epitope is extracellular. We observed that in contrast to wild-type SIDT1, which has a good cell surface localization, SIDT1 mutant was barely detected at the cell surface (Figure 3C). Indeed, in saponin-treated HEK293T cells, SIDT1 mutant was predominantly localized to the endoplasmic reticulum (ER) (Figure 3D and Figure S4). In conclusion, our data indicate that the transport of SIDT1 to the cell surface would be critically impaired in the absence of ECD-localized *N*-glycosylation.

### *N*-glycosylation contributes to RNA binding and protein stability of SIDT1

It is well established that protein *N*-glycan moieties play a crucial role in mediating a wide range of intrinsic and extrinsic interactions (30). There are several examples of the requirement for *N*-glycosylation in the formation of protein-protein interactions. For example, *N*-glycosylation of the vasopressin V1a receptor is required for optimal receptor-ligand binding, and *N*-glycosylation of the P2Y12 receptor is required for proper downstream Gi-mediated signaling (38, 39). Our previous studies and other reports have shown that ECD of the SID-1 transmembrane family can directly bind to RNA, which is a critical activity for its involvement in RNA transport (19, 25). To investigate whether this RNA binding is likely to be *N*-glycosylation dependent, we compared the binding of SIDT1 proteins with different glycosylation status to RNA.

First, we used kifunensine during the SIDT1^ECD^ expression, which is an inhibitor of ER-localized mannosidase I and complex *N*-glycosylation, resulting in the production of glycoproteins with high-mannose oligosaccharides rather than the characteristic complex structures (Figure S5A) (40). We incubated 5’-Carboxyfluorescein (FAM)-labeled RNA with increasing concentrations of purified SIDT1^ECD^, with or without kifunensine treatment, and separated the bound and free forms of RNA on native polyacrylamide gels. After incubation of RNA with SIDT1^ECD^, we observed an unshifted band corresponding to unbound RNA and a compact shifted band presumably corresponding to the saturated RNA-protein complexes. These results showed a progressive decrease in RNA mobility with increasing protein concentration, indicating that SIDT1^ECD^ interacts directly with RNA. Interestingly, we found that no band shifts were observed when kifunensine was applied during SIDT1^ECD^ expression versus wild-type SIDT1^ECD^, suggesting *N*-glycosylation affects the binding of SIDT1^ECD^ to RNA under the above conditions (Figure S5B).

Furthermore, a fusion protein of SIDT1^ECD^ with immunoglobulin fragment crystallizable region (Fc; SIDT1^ECD-Fc^) was generated to enhance the protein stability of SIDT1^ECD^, and deglycosylated SIDT1^ECD-Fc^ was obtained by treatment with PNGase F for analysis of the RNA binding (Figure S5C). EMSA results showed that the interaction of SIDT1^ECD^ with RNA was significantly attenuated when PNGase F was applied (Figure 4A). Full-length SIDT1 with or without PNGase F treatment was analyzed by microscale thermophoresis (MST). Properly glycosylated SIDT1 exhibits high affinity binding to both RNA at pH 5.5, with a dissociation constant (*K*_D_) of 30.97 nM. However, the binding affinity decreased significantly when PNGase F was used, resulting in a *K*_D_ of 6.71 μM (Figure 4B). Collectively, these results suggest that the *N*-glycosylation is critical for SIDT1 binding to RNA, and reduction or absence of *N*-glycosylation reduces SIDT1 binding to RNA. The phospholipase activity of full length SIDT1 was also assessed. Enzymatic activity assays showed that deglycosylated SIDT1 had an essentially unimpaired hydrolytic activity towards short-chain 8:0 phosphatidylcholine (PC), suggesting that *N*-glycosylation is not an absolute requirement for the phospholipase activity of SIDT1 (Figure S5D).

**Figure 4.**
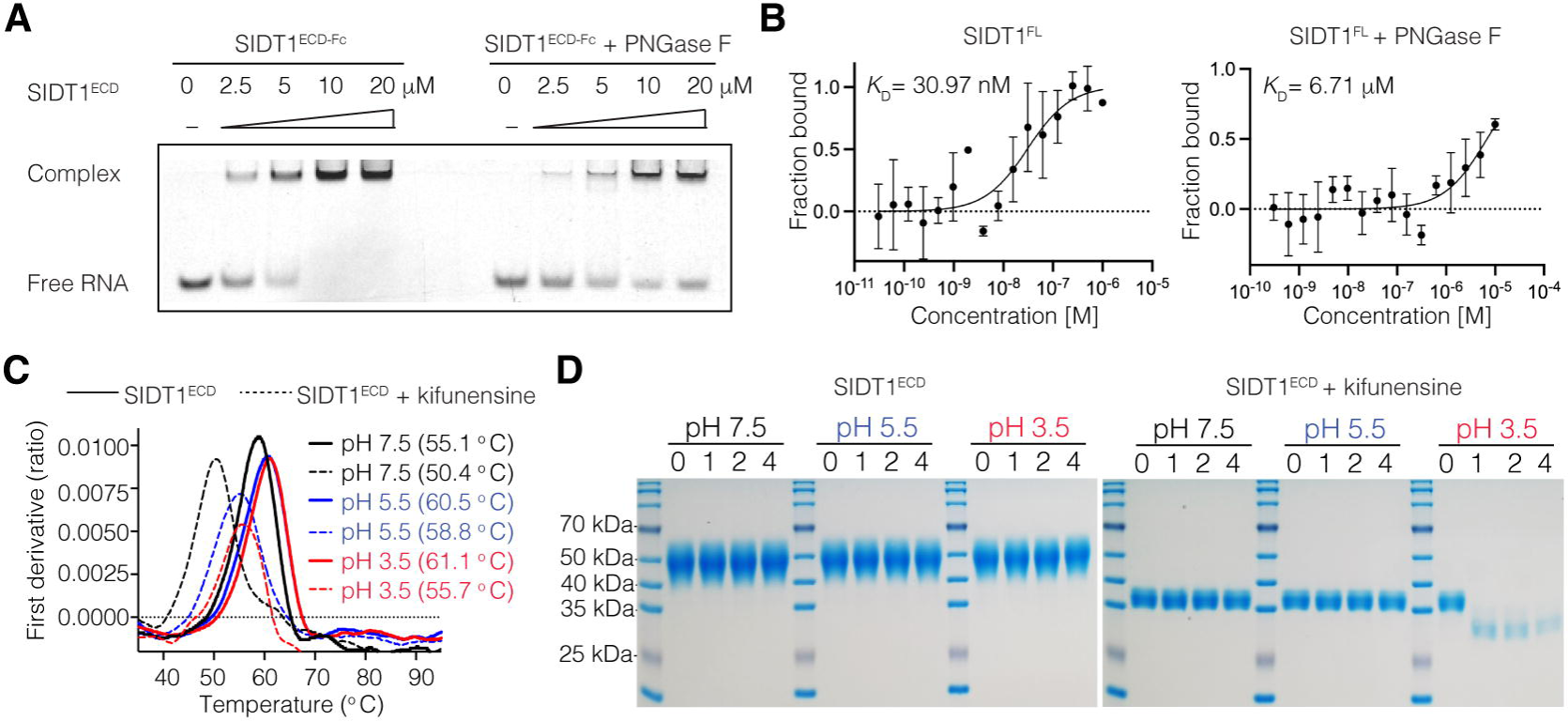
Effect of *N*-glycosylation on RNA binding and protein stability of SIDT1. **A** RNA binding of SIDT1^ECD-Fc^ and PNGase F-treated SIDT1^ECD-Fc^ with RNA as determined by EMSA. The final protein concentrations of SIDT1^ECD-Fc^ in lanes 1-5 are 0, 2.5, 5, 10, and 20 μM, respectively, and the final 5’-FAM-labeled RNA concentration is 2.5 μM. Complex: bound; Free RNA: unbound. **B** MST analysis measuring binding affinities of SIDT1^FL^ (left panel) and PNGase F-treated SIDT1^FL^ (right panel) with RNA at pH 5.5. The calculated dissociation constant (*K*_D_) value represents the affinity between SIDT1^FL^ and RNA. Three independent experiments performed and data are represented as the mean ± S.D. **C** Protein stability of SIDT1^ECD^ (solid line) and kifunensine-treated SIDT1^ECD^ (dotted line) was recorded in real time as the temperature increased from 35 to 95 °C. DSF profiles (ratio between fluorescence at 350 nm and 330 nm) are shown. **D** Time-chase experiments show the degradation of purified SIDT1^ECD^ and kifunensine-treated SIDT1^ECD^ at different pH values (pH 7.5, 5.5, and 3.5; analyzed by SDS-PAGE) and incubation for the experimental period of 0/1/2/4 hours. Three independent experiments were performed. DSF, differential scanning fluorometry; ECD, extracellular domain; EMSA, electrophoretic mobility shift assay; MST, microscale thermophoresis; PNGase F, peptide-N-glycosidase F.

Previous studies have shown that SIDT1 mediates RNA transport in lysosomes and pit cells of the gastric epithelium, where the ECD domain is exposed to the lumen of the lysosome and the acidic gastric environment (11, 18). What’s more, our previous studies have demonstrated the ability of SIDT1 to interact with small RNAs in a pH-dependent manner (19). Since *N*-glycosites are located in the ECD and glycans are generally known to protect against proteolysis (39, 41), we postulate that *N*-glycosylation may function to maintain the protein stability of SIDT1 at low pH and protect it from degradation. Treatment of purified SIDT1^ECD^ with PNGase F resulted in a large amount of precipitation, leaving no glycan-free SIDT1^ECD^ proteins available for further analysis, indicating the important role of *N*-glycans in protein stability. Instead, we examined the protein stability of purified SIDT1^ECD^ and kifunensine-treated SIDT1^ECD^ at different pH values (pH 7.5, 5.5, and 3.5). We measured thermostability as a consequence of protein stability using differential scanning fluorometry (DSF). Temperature-dependent changes in intrinsic protein fluorescence revealed that the two SIDT1 proteins with different *N*-glycosylation states were folded. The *N*-glycosylation deficiency of SIDT1^ECD^ caused a decrease in transition midpoint (Tm) of 4.7 °C at pH 7.5, 1.7 °C at pH 5.5, and 5.4 °C at pH 3.5 compared to SIDT1^ECD^ with a Tm of 55.1 °C, 60.5 °C, and 61.1 °C at pH 7.5, 5.5, and 3.5, respectively (Figure 4C). This indicates that variation in *N*-glycosylation status alters the protein stability of SIDT1.

We then performed a comprehensive time-chase experiment to monitor the degradation of SIDT1^ECD^. We incubated purified SIDT1^ECD^ or kifunensine-treated SIDT1^ECD^ proteins at pH 7.5, 5.5, and 3.5 and then analyzed them on SDS-PAGE gel to visualize the relative mobility. Notably, incubation of SIDT1^ECD^ with low pH buffer for the experimental period of 0, 1, 2, or 4 hours did not result in degradation (Figure 4D). Interestingly, incubation of kifunensine-treated SIDT1^ECD^ at pH 3.5 for the same periods dramatically increased degradation compared to neutral conditions (Figure 4D). The results suggest that the complex *N*-glycosylation protects SIDT1 from degradation in an acidic environment. Taken together, these results indicate that the *N-*glycosylation of SIDT1 enhances its stability and protects it from degradation in an acidic environment.

### Absent *N*-glycosylation can attenuate SIDT1-mediated RNA uptake

Previous studies have shown that the mammalian SID-1 transmembrane family appears to play several cell physiological roles by mediating cellular uptake and intracellular trafficking of oligonucleotides (10, 11, 15, 18, 20, 42). To investigate the potential impact of *N*-glycosylation on SIDT1-mediated RNA transport, a pancreatic ductal adenocarcinoma (PANC1) cell line was used in this study, and the RNA uptake assay was performed as previously described (14). As shown in Fig. 5A, cells were subjected to deglycosylation treatment with PNGase F prior to the RNA uptake assay. Western blotting demonstrated that PNGase F treatment caused a down-shift of the SIDT1 band but did not alter the protein expression level (Figure 5B).

**Figure 5.**
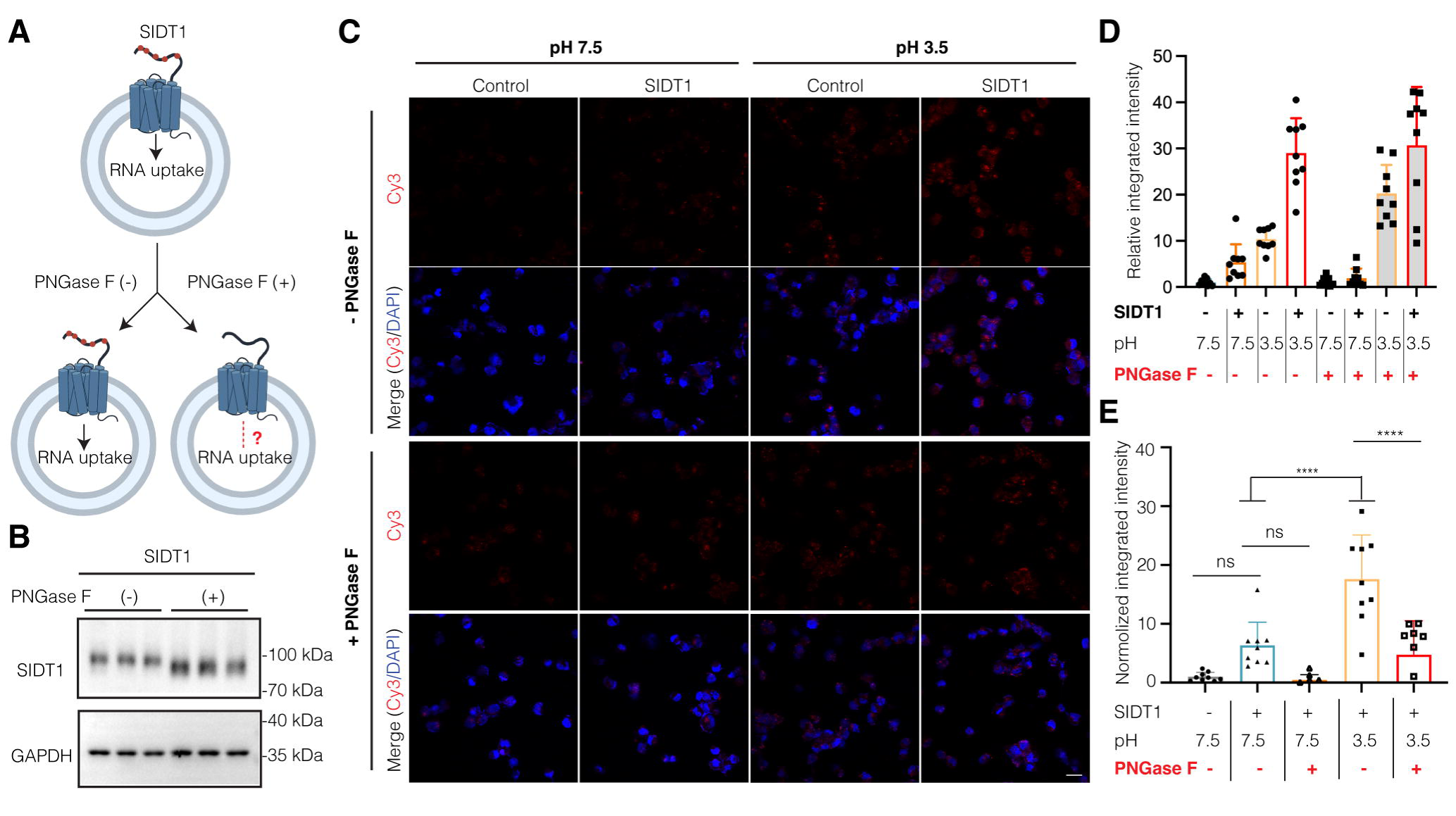
*N*-glycosylation is required for SIDT1-mediated RNA uptake. **A** Schematic diagram of the SIDT1-mediated RNA uptake assay. pcDNA 3.1(+) and SIDT1 transfected cells were treated with or without PNGase F for 1 h before the RNA uptake assay. **B** Immunoblots of representative PNGase F-treated and -untreated PANC1 cells. The cell suspension samples were immunoblotted for SIDT1 (upper) and GAPDH (lower). GAPDH served as a loading control. Three independent experiments were performed. **C** Representative images of PNGase F-treated (+) and untreated (−) PANC1 cells after the uptake of synthetic Cy3-labeled RNA (red) and DAPI (blue) was used for nuclear staining, Scale bar, 20 μm. **D, E** The statistical analyses of PNGase F-untreated (−) and treated (+) PANC1 cells after the uptake of synthetic Cy3-labeled RNA, corresponding to figure 5C. The raw data (**D**) and normalized (**E**) analyses are shown (n=9). Data are represented as the mean ± S.D. The statistical significance is determined by One-way ANOVA. ns, not significant, ***P < 0.001, ****P < 0.0001. DAPI, 2-(4-amidinophenyl)-6-indolecarbamidine dihydrochloride; PANC1, pancreatic ductal adenocarcinoma; PNGase F, peptide-N-glycosidase F.

The PANC1 cells were transfected with empty vector pcDNA3.1(+) and wild-type SIDT1, respectively. After incubation with synthesized Cy3-labeled RNA, RNA uptake was detected by fluorescence (Figure 5C). The Cell Counting Kit-8 (CCK-8) assay demonstrated that PNGase F treatment for 60 minutes and at pH 3.5 for 10 minutes did not impair the cell viability of PANC1 cells (Figure S6). All data were statistically analyzed and background (PNGase F-untreated cells) deductions were applied to normalize the data for PNGase F-treated cells (Figure 5D and 5E). We observed that incubation of cells with RNA in medium at pH 3.5 for 10 minutes dramatically increased RNA uptake in wild-type SIDT1-transfected cells than that of RNA at pH 7.4 in cultured cells, which is in agreement with previous studies that SIDT1-mediated RNA uptake is dramatically enhanced in a pH-dependent manner (Figure 5C-E) (18). At pH 7.5, the RNA uptake capacity of deglycosylated SIDT1 was slightly attenuated compared to that of cells expressing wild-type SIDT1 (Figure 5C-E). In contrast, PNGase F-treated cells, which have similar expression levels to the wild-type SIDT1, showed a significantly reduction in RNA uptake at pH 3.5 (Figure 5C-E). Our results show that cells expressing wild-type SIDT1 exhibited enhanced RNA uptake compared to PNGase F-treated cells, suggesting that *N*-glycosylation, to some extent, plays a critical role in the SIDT1-mediated RNA uptake.

In conclusion, our data indicate that ECD-localized *N*-glycosylation is important for the RNA binding and SIDT1-mediated RNA uptake. Removal of the *N*-glycans resulted in decreased RNA binding of SIDT1 and subsequently severe defects in RNA uptake activity. What’s more, our results indicate that *N-*glycosylation of SIDT1 enhances its stability and protects it from degradation in an acidic environment, which may be one of the reasons for the reduced RNA binding and uptake.

## Discussion

Protein *N*-glycosylation is a post-translational modification and can significantly affect the properties and functions of target proteins by the addition of bulky, hydrophilic carbohydrate structures to specific asparagine residues (30). *N*-glycosylation has previously been shown to regulate protein folding, solubility, stability and activity, protection from proteases, subcellular targeting, and other interactions (33, 36). Our sequence analysis revealed that *N*-glycosylation of SID1 transmembrane family members is conserved in most mammals, suggesting potentially functional contributions of *N*-glycosylation to the mammalian SID-1 transmembrane family. In this study, we analyzed the role of *N*-glycosylation in SIDT1-mediated RNA uptake function and explored the underlying molecular mechanisms. Thus, the evidence provides a direct link from *N*-glycosylation to SIDT1-mediated RNA uptake.

In agreement with previous studies, we verified that SIDT1 and SIDT2 are *N*-linked glycosylated by glycosidase digestion (PNGase F). We profiled the site-specific *N*-glycosylation of recombinant SIDT1 and SIDT2 proteins expressed from HEK293F cells. The occupancy at ECD-located *N*-glycosites of SIDT1 and SIDT2 (N57, N67, N83, N136, N282 for SIDT1; N27, N54, N60, N123, N141, N165 for SIDT2) was quantified. The results show that the putative *N*-glycosites between the TM domains are not occupied. It is likely that the proximal putative *N*-glycosites could not be recognized by the oligosaccharyltransferase, since a minimum of 12 amino acid residues between the *N*-glycosylation consensus site and the transmembrane helix is required for efficient *N*-glycosylation (43).

Our results show that the occupancy of glycosylation at each *N*-glycosite is different, and the *N*-glycosylation of SIDT1 and SIDT2 in human cells displays remarkable heterogeneity, with paucimannose, high mannose, hybrid and complex-type glycans present across the *N*-glycosites. Various studies have shown that protein *N*-glycosylation patterns can change in disease. In non-alcoholic fatty liver disease (NAFLD) and non-alcoholic steatohepatitis (NASH), the glycosylation status of SLC transporters is altered (44). The expression of non-glycosylated organic anion transporting polypeptides OATP1B1/3, OATP2B1 and non-glycosylated Na^+^-taurocholate co-transporting polypeptide (NTCP) were found to be increased in NASH using gene array data and subsequent Western blot analysis of human liver donors (45). In several diseases, SIDT1 and SIDT2 are implicated. SIDT1 plays a key role in breast, pancreatic, and non-small cell lung cancer (7-9). Loss of function studies has demonstrated an essential role for SIDT2 in the regulation of glucose metabolism, lipid metabolism, antiviral immunity, skeletal muscle homeostasis, and tumor development (5-8, 12, 13). It is also implicated in neurodegenerative disorders like Alzheimer’s (46). It would therefore be interesting to determine whether *N*-glycosylation is altered in SIDT1- and SIDT2-related diseases.

A large body of evidence highlights an important role for *N*-glycosylation in the cell surface expression of biomolecules (35, 41, 47). Our results show that complete deglycosylation of SIDT1 by site-directed mutagenesis results in a significant decrease in cell surface expression of the SIDT1, together with increased retention in the intracellular compartment. Colocalization studies showed that non-*N*-glycosylated SIDT1 was localized predominantly to the ER and not to the Golgi apparatus, suggesting that *N*-glycosylation is important for the biosynthetic processing in the ER. In the absence of *N*-glycans, processing in the Golgi apparatus is prevented, thereby blocking the Golgi secretory pathway. Similar results were observed for the secretory expression of SIDT1^ECD^, showing that non-*N*-glycosylated variant has a significant and substantial secretion defect. These results correlated with the transcriptional up-regulation of molecules HYOU1, HSPA5, etc, indicating cells undergoing ER stress and activation of pathways within the unfolded protein response. Our results indicate that *N*-glycosylation is required for efficient protein expression of SIDT1. However, the exact role of *N*-glycosylation, particularly the impact of different *N*-glycosites on SIDT1, still requires further investigation to be elucidated.

Another potential role of *N*-glycosylation is to affect the interaction between SIDT1 and RNA. Here, we show that the complete removal of *N*-glycans disrupts the interaction of SIDT1 with RNA, suggesting that *N*-glycans play a critical role in shaping the SIDT1-RNA interaction. A likely explanation for the reduced RNA binding is that the *N*-glycans are involved in the binding of SIDT1 to RNA. Further single-molecule studies are needed to determine the detailed biochemical and biophysical properties of SIDT1 involved in RNA binding. Another plausible explanation is the reduced protein stability of *N*-glycan-deficient SIDT1. Previous studies have clearly shown that dietary miRNAs are absorbed occurs in the stomach, and that gastrically enriched SIDT1 is the key molecule (18). In the gastrointestinal tract, SIDT1 has the highest expression levels in the stomach and large intestine (including the cecum, colon and rectum), and these are the tissues exposed to the highest proteolytic activities in the body with various proteases derived from pancreatic secretion. ECD of SIDT1 in the apical membrane are therefore particularly susceptible to rapid degradation with loss of functionality. Indeed, our results suggest that *N*-glycosylation of SIDT1 confers resistance to the mimicked acidic conditions (pH 3.5).

In our RNA uptake assay, we cannot rule out the possibility that acidic culture or PNGase F-treated conditions *in vitro* might influence all cell surface proteins in cells, leading to some indirect effects. However, the current study suggests that *N*-glycosylation is to some extent required for efficient SIDT1-mediated RNA uptake, which may be attributed to the fact that *N*-glycosylation is required for correct subcellular localization of SIDT1, efficient SIDT1-RNA interactions, and sufficient protein stability. It would be interesting to further study whether the reduced RNA uptake is due to the aberrant glycan structure of SIDT1 or to the specific *N*-glycosite.

Over the past few decades, RNA-based therapies have developed rapidly compared to conventional drugs such as small molecules and proteins, and have many advantages, such as easier and faster design and production, cost-effectiveness, an expanded range of therapeutic targets, and flexible combinations of drug cocktails for personalized treatment (48-50). RNA-based therapeutics, such as siRNA, miRNA, and mRNA, can silence gene expression, regulate target genes, or induce the expression of specific genes (51, 52). Exogenous dietary miRNAs can be transported to the pit cells in the stomach by the intrinsic carrier protein SIDT1 and then secreted as functional entities in exosomes (18). This natural mammalian uptake pathway of dietary miRNAs can be easily exploited for oral delivery of therapeutic miRNAs, which may be an important direction for the future development of RNA-based medicine. Our findings reveal the effect of *N*-glycosylation on SIDT1-mediated RNA uptake and shed light on the development of oral delivery-based small RNA therapeutics.

## Experimental Procedures

### Plasmid construction

The full-length cDNAs for human SIDT1 and human SIDT2 were synthesized by Genscript Company (SIDT1, Uniprot: Q9NXL6; SIDT2, Uniprot: Q8NBJ9). For protein expression and purification, the sequence encoding residues 23-310 of human SIDT1 and residues 22-292 of human SIDT2 were cloned into a modified pcDNA3.1(+) vector with a melittin signal sequence (MKFLVNVALVFMVVYISYIYA) at the N-terminus and a 6 x His tag at the C-terminus. To ensure the dimerization and stability, we utilized a fusion strategy by adding an Fc fragment to the C-terminus of the ECD to facilitate dimerization as our previous study (19). The final construct was subcloned into the pMlink expression vector, and clone identities were verified by Sanger sequencing. Full-length SIDT1 and SIDT2 with a C-terminal Flag tag and a N-terminal efficient signal peptide derived from the influenza virus hemagglutinin HA (MKANLLVLLCALAAADA) or myc tag were subcloned into pcDNA3.1(+) vector. The SIDT1 mutants were generated by PCR-based mutagenesis method and cloned into this vector in the same way. The identity of each clone was verified by Sanger sequencing.

### Protein expression and purification

SIDT1^ECD^ and SIDT2^ECD^ expression were performed in HEK293F cells (Invitrogen). Cells were cultured in OPM-293 CD05 medium (OPM Biosciences co., Ltd.) at 37 °C supplemented with 5% CO_2_ in a ZCZY-CS8 shaker (Shanghai Zhichu Instrument co., Ltd.) at 120 rpm. When the cell density reached 2.5 × 10^6^ to 3.0 × 10^6^ cells per mL, the cells were transiently transfected with the expression plasmids. Approximately 1.5 mg of plasmids were pre-mixed with 4.5 mg of polyethylenimines (PEIs) (Polysciences) in 50 mL of fresh medium for 30 min before application. For transfection, 50 mL of the mixture was added to 1 liter of cell culture, and the transfected cells were harvested 72 hours after incubation. Cells were collected by double centrifugation and 6x His-tagged target protein in the supernatant was purified using the medium by Ni-NTA (Qiagen) affinity chromatography. Protein was eluted with buffer B containing 50 mM HEPES pH 7.5, 250 mM NaCl, and 250 mM imidazole. The eluate was then concentrated and purified by high resolution gel filtration column Superdex 200 Increase 10/300 (GE Healthcare), pre-equilibrated with buffer C containing 20 mM HEPES pH 7.5, 100 mM NaCl and 0.1 mM tris (2-carboxyethyl) phosphine hydrochloride (TCEP). Peak fractions of the targets were confirmed by SDS-PAGE followed by Coomassie blue staining, and data analysis were performed using GraphPad Prism 9.0 software (GraphPad Software, San Diego, CA, USA).

Full-length SIDT1 and SIDT2 expression were performed in HEK293F cells in the same manner. For protein expression and purification, 2 liters of cells were harvested 60 hours post-transfection, then the cells were lysed in lysis buffer containing 50 mM HEPES pH 7.5, 300 mM NaCl, and supplemented with protease inhibitor cocktail (MCE, Cat# HY-K0010) and 1 mM phenylmethylsulfonyl fluoride (PMSF). Cells were disrupted by continuous high-pressure cell disruption at 800 bars. Cells debris was removed by low-speed centrifugation for 30 minutes and the supernatant were further subjected to high-speed centrifugation at 150,000 g for 90 minutes using a Ti-45 rotor (Beckman), and the membrane component was collected. The membrane was resuspended and homogenized using a glass dounce homogenizer for 20 cycles on ice in solubilization buffer containing 50 mM HEPES pH 7.5, 300 mM NaCl, 1 mM EDTA, 2% (*w/v*) n-dodecyl-β-D-maltoside (DDM, Anatrace), 0.2% (*w/v*) cholesteryl hemisuccinate (CHS, Anatrace), protease inhibitor cocktail and solubilized at 4 °C with gentle agitation for 2 hours. The extraction was centrifuged at 39,000 g (Beckman) for 1 hour to remove the insoluble components, and supernatant was applied to incubate with anti-Flag affinity resin (Genscript) at 4 °C for 1.5 hours. The protein was eluted using buffer containing 50 mM HEPES pH 7.5, 300 mM NaCl, 0.01% (*w/v*) lauryl maltose neopentyl glycol (LMNG), 0.001% (*w/v*) CHS, and 500 μg/mL synthesized Flag peptide (Genscript). The eluent was further purified by SEC using a Superose 6 increase 10/300 column (GE Healthcare) pre-equilibrated with buffer containing 25 mM HEPES pH 7.5, 150 mM NaCl, 0.5 mM EDTA, 0.01% (*w/v*) LMNG, and 0.001% (*w/v*) CHS. Protein purity was examined by SDS-PAGE and visualized by Coomassie blue staining. Peak fractions were pooled and concentrated to 2 mg/mL, and then flash frozen in liquid nitrogen for LC/MS analysis.

The plasmid encoding PNGase F was gifted from the laboratory of Fuquan Yang (Institute of Biophysics, Chinese Academy of Sciences) and a C-terminal Strep II tag was designed for purification. The plasmid was transformed into *E. coli* BL21 (DE3) cells. Cells were grown to an OD_600_ of 0.8 at 37 °C and then induced with 0.2 mM isopropyl-1-thio-D-galactopyranoside (IPTG). The cells were harvested by centrifugation after shaking at 16 °C for 20 hours and resuspended in buffer containing 50 mM Tris-HCl pH 8.0, 500 mM NaCl, and 1 mM PMSF, followed by disruption with a high-pressure homogenizer cell disrupter (JNBIO), and the lysate was centrifuged for 30 minutes at 18,000 g. The supernatant was loaded onto Strep-Tactin affinity beads (GE Healthcare) and eluted in buffer containing 50 mM Tris-HCl pH 8.0, 300 mM NaCl, and 2.5 mM desthiobiotin. The protein was subsequently loaded onto a Superdex 200 increase 10/300 column pre-equilibrated with buffer containing 20 mM Tris-HCl pH 8.0, 150 mM NaCl, and 0.5 mM TCEP. Purified proteins were assessed by SDS-PAGE and stored at -80 °C.

### Transfection and Western blot

Transfections of cell monolayers were performed using the Lipofectamine 3000 (Invitrogen) in 6-well plates according to the manufacturer’s instructions. Transfected cells were incubated at 37°C for 24 hours unless otherwise stated. To obtain cell lysates, cells were trypsinized, washed with phosphate-buffered saline (PBS), and lysed in lysis buffer with 50 mM Tris-HCl pH 7.5, 150 mM NaCl, 1 mM EDTA, 1.5 mM MgCl_2_, 1% NP-40, 1.5 mM PMSF, and protease inhibitor cocktail (MCE, Cat# HY-K0010). Lysates were cleared by centrifugation at 14000 g and 4 °C for 10 minutes. The total protein in the supernatant was quantified using the bicinchoninic acid assay (BCA) kit (Beijing TianGen Biotechnology Co., Ltd.). For each sample, equivalent amounts of total protein were evaluated during the analysis. Cell lysates or media were denatured by boiling in 4 x SDS-PAGE sample loading buffer (200 mM Tris-HCl pH 6.8, 40% (*v/v*) glycerol, 8% (*w/v*) SDS, 0.008% (*w/v*) bromophenol blue, and 20% (*v/v*) β-mercaptoethanol), separated on 10% or 12.5% SDS-PAGE gels, and transferred to PVDF membranes. Blots were imaged after incubation with appropriate primary and secondary antibodies using the Tanon-5100 Fluorescent Imaging System (Tanon Science & Technology), followed by quantification as required using ImageJ as appropriate.

### De-*N*-glycosylation analysis

Whole cell lysates (WCLs) and culture medium were obtained from HEK293T cells transfected with SIDT1 or SIDT2. The combinations were subjected to de-*N*-glycosylation analysis to confirm the extent of glycosylation using PNGase F, which cleaves the innermost *N*-acetyl-glucosamine (GlcNAc) and asparagine residues (Asn, N) residues. The samples were treated with PNGase F at an enzyme/protein ratio of 1:50 (*w/w*) at 37 °C for 3 hours. The reaction mixture was examined on 10% SDS-PAGE to check the degree of glycosylation of the recombinant proteins. To obtained deglycosylated SIDT1^ECD^ and full length SIDT1, purified recombinant protein samples were treated with PNGase F at an enzyme/protein ratio of 1:50 (*w/w*) at 37 °C for 1 hours.

### In-solution digestion of samples

The purified samples were digested into peptides as previously described (53). Briefly, samples were reduced with DTT at a final concentration of 10 mM at 30 °C for 1 hour. The resulting free thiols were alkylated with IAM at a final concentration of 40 mM for 45 minutes at room temperature (RT) in the dark. The same amount of DTT was then added to remove excess IAM at 30 °C for 30 minutes. Proteins were then digested with Lys-C (Wako Pure Chemical Industries, Osaka, Japan) at an enzyme-to-protein ratio of 1: 100 (*w/w*) at 37 °C for 3 hours. After dilution with 50 mM Tris-HCl pH 8.0, samples were digested with sequencing grade modified trypsin (Promega, Madison, WI, USA) at an enzyme-to-protein ratio of 1:50 (*w/w*) at 37 °C overnight. Enzymatic digestion was stopped with formic acid (FA) and the supernatant was collected after centrifugation at 20,000 × g for 20 minutes. Peptides were desalted on HLB cartridges (Waters, Milford, MA, USA) and dried in a benchtop centrifugal vacuum concentrator (Labconco, MO, USA).

### LC-MS/MS analysis and data analysis

Peptide samples were dissolved in 0.1% FA and analyzed on the Orbitrap Eclipse Tribrid Mass spectrometer (Thermo Fisher Scientific) coupled to an Easy-nLC 1200 HPLC system (Thermo Fisher Scientific). Samples were separated on a fused silica trap column (100 μm ID×2 cm) in-house packed with reversed-phase silica (Reprosil-Pur C18 AQ, 5 μm, Dr. Maisch GmbH, Baden-Wuerttemberg, Germany) coupled to an analytical column (75 μm ID×20 cm) packed with reversed-phase silica (Reprosil-Pur C18 AQ, 3 μm, Dr. Maisch GmbH). The peptides were analyzed with 103 min gradient (buffer A: 0.1% FA in H_2_O, buffer B: 80% ACN, 0.1% FA in H_2_O) at a flow rate of 300 nL/minutes (4-11% B, 4 min; 11-22% B, 32 minutes; 22-32% B, 31 minutes; 32-42% B, 23 minutes; 42-95% B, 3 minutes; 95% B, 10 minutes). MS data were acquired in data-dependent acquisition mode. The cycle time was set as 3 seconds (s). The spray voltage of the nano-electrospray ion source was 2.0 kV, with no sheath gas flow, the heated capillary temperature was 320 °C. An MS1 scan was acquired from 350 to 2000 m/z (60,000 resolution, 4e^5^ AGC, normalized AGC=100%, maximum injection time=50 ms) followed by EThcD MS/MS acquisition of the precursors in an order of intensity and detection in the Orbitrap (30, 000 resolution, 4e5 AGC, normalized AGC=800%, maximum injection time=200 ms).

All raw data files were searched against the UniProt human reviewed protein database with target protein sequences and common contaminants, using the PTMcentric search engine Byonic (version 3.11.3, Protein Metrics) (54) incorporated in Proteome Discoverer (PD 2.2). Trypsin was selected as the enzyme and two maximum missed cleavages were allowed. Searches were performed with a precursor mass tolerance of 10 ppm and a fragment mass tolerance of 0.02 Da. Static modifications consisted of carbamidomethylation of cysteine residues (+57.02146 Da). Dynamic modifications consisted of the oxidation of methionine residues (+15.99492 Da), deamidation of asparagine and glutamine (+0.98402 Da), and *N*-glycosylation on asparagine. Oxidation and deamidation were set as “rare” modifications, and *N*-glycosylation was set as a “common” modification through Byonic node. Two rare modifications and one common modification were allowed. 182 human *N*-glycans embedded in Byonic was used. Results were filtered to a 1% protein FDR as set in the Byonic parameters, and data were further processed to 1% FDR at the PSM level using the 2D-FDR score (a simple variation of the standard target-decoy strategy that estimates and controls PSM and protein FDRs simultaneously) (55). Only those *N*-glycopeptides with Byonic score >100 and |logProb| > 1 were reported. Each glycopeptide identified should have the consensus motif N-X-S/T, X≠P. The filtering criteria have been reported to result in confident glycosite assignment at glycopeptide spectral match level (56).

### Electrophoretic mobility shift assay

The interaction between SIDT1^ECD^, kifunensine-treated, or PNGase F-treated SIDT1^ECD^ and RNA were examined by EMSA. The RNA (5’-UCGCUUGGUGCAGAUCGGGAC-3’) labeled at the 5’ end with FAM was synthesized by Genscript co., Ltd, and dsRNA was obtained by annealing with an equimolar amount of the reverse strand. For the EMSA, increasing amounts of the SIDT1^ECD^ proteins (1.25, 2.5, 5, and 10 μM) were incubated with 2.5 μM FAM-labeled ssRNA in a reaction EMSA buffer containing 40 mM Tris-HCl pH 5.5, (adjusted by acetate), 100 mM NaCl, 2.5 mM EDTA and 5 % (*v/v*) glycerol to a total volume of 20 μl at RT for 30 minutes. The obtained products were then resolved on 5 % native acrylamide gels (37.5:1 for acrylamide: bisacrylamide) in 1x Tris-acetate-EDTA (TAE) running buffer (pH 5.5) under an electric field of 10 V/cm for about 20 minutes on ice. The gel was visualized and analyzed by the Tanon-5100 Fluorescent Imaging System and Image J software.

### Microscale thermophoresis

The MST experiments were conducted using a Monolith NT.115 instrument (NanoTemper Technologies). Purified SIDT1^FL^ and PNGase F-treated SIDT1^FL^ were exchanged into a labeling buffer containing 25 mM NaHCO_3_ pH 8.3, 100 mM NaCl, 0.05% Tween-20, 0.01% (*w/v*) LMNG, and 0.001% (*w/v*) CHS. After buffer exchange, the proteins were labeled with a RED-NHS Labeling Kit (NanoTemper Technologies) according to the manufacturer’s instructions. For the MST measurements, the labeled proteins were dialyzed and exposed to pH 5.5 conditions buffer (25 mM MES, 100 mM NaCl, and 0.05% Tween-20). Throughout the experiments, the labeled protein concentration was maintained at 5 nM. 16-step serial dilutions of RNA were prepared in the same buffer to ensure consistent buffer conditions. MST measurements were performed at a constant temperature of 25 °C, with a 5-second LED on-time followed by a 30-second MST on-time. The LED and MST power settings were optimized for each experiment to achieve optimal signal-to-noise ratios and minimize aggregation or adsorption effects. Data were collected and analyzed using the MO. Affinity Analysis software (NanoTemper Technologies). To ensure the accuracy and reliability of the results, MST experiments were performed in triplicate, and the data were averaged to determine the final *K*_D_ values.

### *In vitro* phospholipase activity assay

The *in vitro* phospholipase activity assay was performed using the Amplex Red Phospholipase D Assay Kit (A12219, Invitrogen) and 1,2-dioctanoyl-sn-glycero-3-phosphocholine (8:0 PC) (Cat# 850315C, Avanti Polar Lipids) as the phospholipid substrate. A positive control sample was prepared with a known concentration of 10 μM H_2_0_2_. The phospholipase activity detection was evaluated using the previously reported method (27). Briefly, 4 μM of purified SIDT1 or PNGase F-treated SIDT1 proteins were incubated with 1 mM PC lipid at 37L for 60 minutes. The reaction mixture was composed of 25 mM HEPES at pH 7.5, 150 mM NaCl, 1 mM PC, 0.01% (*w/v*) LMNG, and 0.001% (*w/v*) CHS, with a total volume of 20 μL. The reaction was terminated by heating at 90°C for 10 minutes, followed by cooling on ice for 20 minutes. Then, a 20 μL working solution was added to the reaction mixture. The working solution contained 50 mM Tris-HCl at pH 8.0, 100 µM Amplex Red, 2 U/mL horseradish peroxidase, and 0.2 U/mL choline oxidase. The reaction mixture was incubated at 37°C for 30 minutes in the absence of light. Then, 40 μL of 50 mM HEPES pH 7.5 was added. Fluorescence measurements were taken using a SpectraMax M4 Microplate Reader (Molecular Devices) in 96-well plates with 540 nm excitation and 590 nm emission wavelengths. Prism 8 software (GraphPad) was used to perform statistical analysis and determine significant differences between samples and controls.

### DSF

The thermal stability was performed using Tycho NT.6 (NanoTemper Technologies). SIDT1^ECD^ and kifunensine-treated SIDT1^ECD^ were diluted to a final concentration of 0.25 mg/mL at various pH (pH 7.5, 5.5, and 3.5), and analyzed in triplicates in capillary tubes. Protein unfolding was monitored using the intrinsic fluorescence at 350 and 330 nm with temperature gradient from 35 to 95 °C at a rate of 30 K/minutes. Data analysis, data smoothing, and the calculation of derivatives involved using the internal evaluation features of the Tycho instrument.

### Time-course experiments

To assess the effect of *N*-glycosylation on the protein stability of SIDT1, SIDT1^ECD^ and kifunensine-treated SIDT1^ECD^ were exposed to the various pH buffer (pH 7.5, 5.5, and 3.5). The reaction was stopped after 0/1/2/4 hours incubation at RT by adding 4 x SDS-PAGE sample buffer. Proteins were heated at 95 °C for 10 minutes and separated by SDS-PAGE.

### Cell Counting Kit-8 assay

PANC1 cells (Invitrogen) were plated in 96-well plates at the density of 8000 cells/well and cultured overnight. On the next day, their medium was replaced with fresh medium supplemented with PNGase F, or the pH of the culture medium was adjusted with hydrochloric acid to 3.5 (100 μL/well). After incubation, 10 μL of the cell counting reagent CCK-8 cell counting kit (Vazyme, Nanjing, China) was added to each well and the cells were incubated for 2 h at 37 °C in 5% CO_2_. After incubation, the optical density (OD) value of each well was measured at 450 nm. Three independent experiments were performed.

### Cell-based RNA uptake assays

Cell-based RNA uptake assays were performed in cultured cells using a previously described method (18, 42). Briefly, equal amounts of the PANC1 cells were seeded on glass coverslips pretreated with TC (Solarbio) in 6-well plates. After culturing for 12 h to reach 60-70% confluency, the cells were transfected with pcDNA3.1(+) or SIDT1 plasmids using Lipofectamine 3000 (Invitrogen). After 24 hours, PANC1 cells were washed twice with 1 mL PBS and treated with or without PNGase F at 37 °C for 1 h. RNA uptake was performed at RT for 10 minutes by incubating cells with 160 μM Cy3-labeled RNA (encoding sequence 5’ -TCGCTTGGTGCAGATCGGGAC-3’) in 1 mL fetal bovine serum (FBS)-free DMEM, or the pH of the culture medium was adjusted with hydrochloric acid to 3.5. The reaction was stopped by three washes with PBS and incubated with FBS-free medium containing 0.2 mg/mL RNase A (Thermo Fisher Scientific, Cat# 12091021) for 1 hours to digest the extracellularly attached RNAs. Cells were washed three times, and total transported RNA and protein expression were detected.

### Immunofluorescence microscopy

Monolayers of HEK293T (Invitrogen) or PANC1 cells grown on coverslips were transfected as described above. After 24 hours of transfection, culture media was removed and the coverslips were washed with PBS. The cells were then fixed with 4% paraformaldehyde at RT for 10 minutes and then incubated with 0.5% saponin for 10 minutes to permeabilize the cells. Coverslips were then incubated in a blocking solution of 3% BSA at RT for 1 hours. Cells were labeled overnight with the anti-Flag, or anti-myc antibody (Abclonal; 1:500) in 3 % BSA at 4 °C, followed by either the anti-PDI (Beyotime Biotechnology; 1:200) or anti-GM130 antibody (Beyotime Biotechnology; 1:200). Between antibodies, the coverslips were washed three times with PBS. The secondary antibody (Alexa Fluor 647-conjugated anti-mouse and Alexa Fluor 488-conjugated anti-rabbit, Abclonal; 1:500) were used to incubate cells at RT in darkness for 1 hours. Both secondary antibodies, Alexa Fluor 647-conjugated anti-mouse and Alexa Fluor 488-conjugated anti-rabbit (Abclonal; 1:500) were applied simultaneously in the 3% BSA solution for 1 hours at RT. The coverslips were then washed three times with PBS and labeled with 2-(4-Amidinophenyl)-6-indolecarbamidine dihydrochloride (DAPI) (Beyotime Biotechnology) for 10 minutes at RT. Images were obtained with a Zeiss LSM 980 scanning confocal microscope. The images were processed with ZEN software (Carl Zeiss).

### Quantification and statistical analysis

Measurements were performed in ImageJ (National Institutes of Health). Quantification and data analysis were performed with GraphPad Prism. The statistical details of experiments can be found in figure legends. All experiments were repeated independently for more than three times. All statistic values were displayed as means ± SD, the statistical significance in this study is determined by One-way analysis of variance (ANOVA) followed by Tukey’s correction or the unpaired, two-tailed Student’s *t*-test, p > 0.05 is set to not significant, *p < 0.05, **p < 0.01, ***p < 0.001, ****p < 0.0001.

### Data availability

The mass spectrometric data have been deposited to the ProteomeXchange Consortium (http://proteomecentral.proteomexchange.org) via the iProX partner repository (57, 58) with the dataset identifier PXD043824. The link access to the raw data is https://proteomecentral.proteomexchange.org/cgi/GetDataset?ID=PXD043824. The data that support this study are available within the article and its Supplementary Information files. All mRNA-seq datasets generated for this study have been deposited at GEO under the accession number GSE239877. The link access to the raw data is https://www.ncbi.nlm.nih.gov/geo/query/acc.cgi?acc=GSE239877.

## Acknowledgements

We thank Qipeng Zhang and Jiatong Tang for valuable assistance and discussion of RNA uptake assay, and Shoubing Zhan for helpful assistance of RT-PCR. We also thank Tongyang Zhu and the National Key Laboratory of Pharmaceutical Biotechnology at Nanjing University in the technical assistance. We are grateful to Jifeng Wang from the Laboratory of Proteomics, Institute of Biophysics, Chinese Academy of Sciences for his help with mass spectrometric data acquisition. This work was supported by funds as following dataset.

## Funding

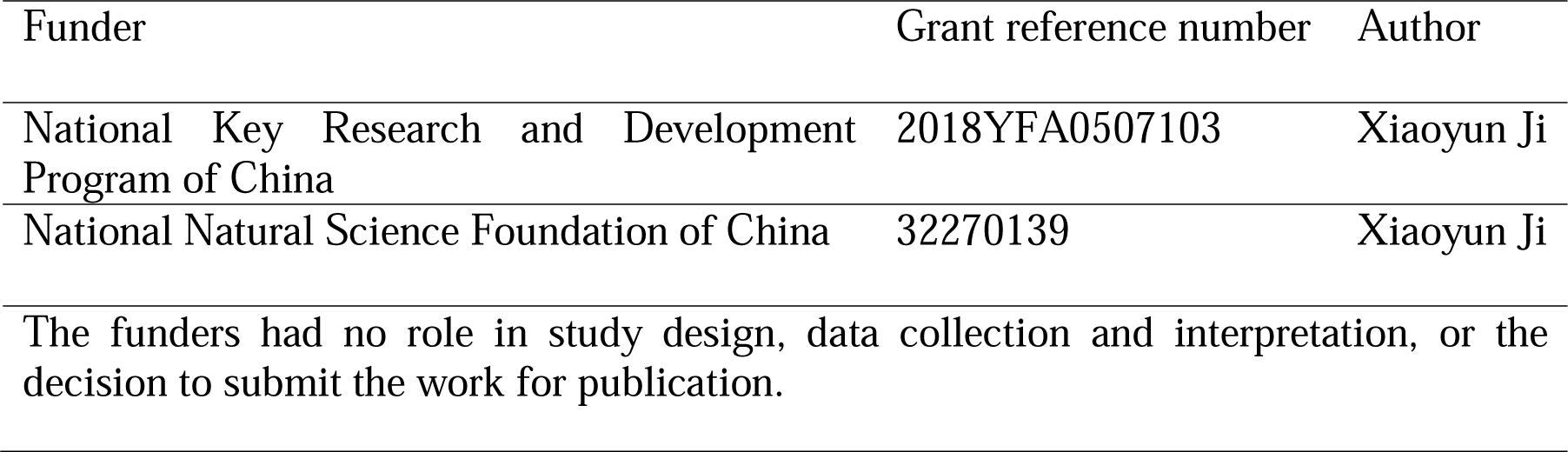

## Corresponding authors

Correspondence to Chenyu Zhang, Fuquan Yang, Xiaoyun Ji.

## Additional information

This article contains supporting information (59-62).

