## Supporting Information for "Characterization of *N*-glycosylation and its Functional Role in SIDT1-Mediated RNA Uptake"

**Supplementary figures S1-S6**

**Supplementary Table S1 (separate excel file): Analysis related to Figure 1D.**

**Supplementary Table S2 (separate excel file): Analysis related to Figure 1E.**

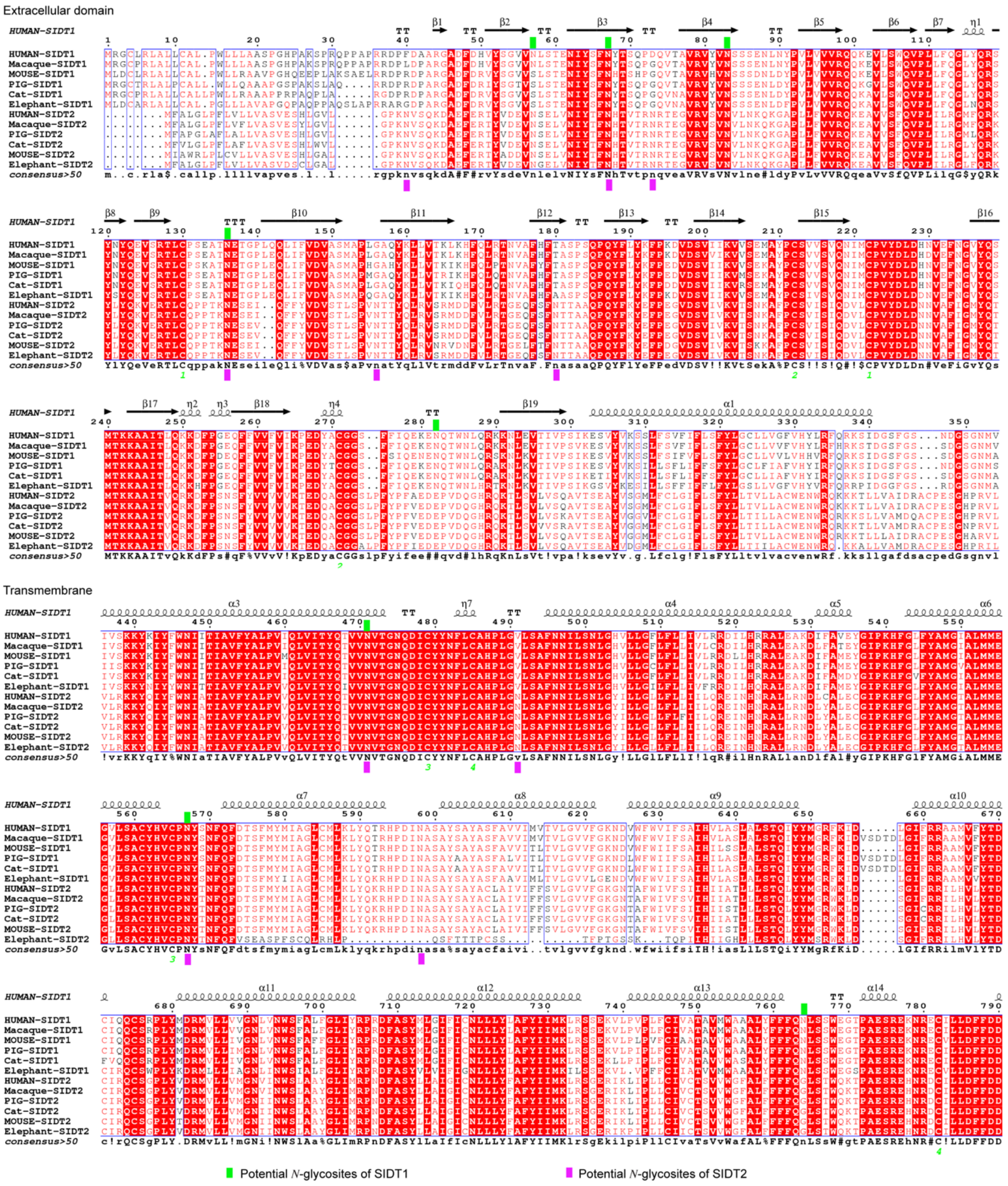


**Figure S1. Potential *N*-glycosites of human SIDT1 and SIDT2.** Sequence alignment of the representative sequences across different mammalian species, including human, macaque, mouse, pig, cat, and elephant. Secondary structure elements predicted by AlphaFold are indicated by arrows for β strands and cylinders for α helices(1,2). Potential *N*-glycosites aligned to human SIDT1 and SIDT2 are highlighted in green and purple, respectively. Sequences were aligned using Clustal Omega(3) and visualized using ESPript 3.0(4).

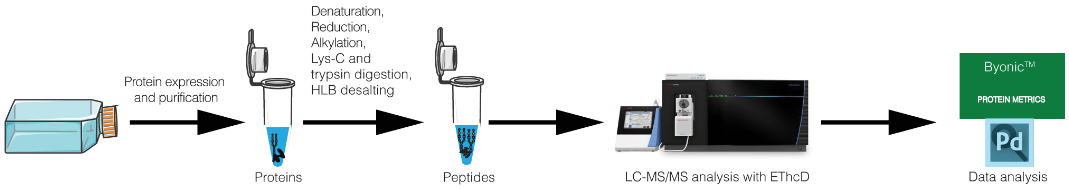


**Figure S2. The workflow for site-specific *N*-glycosylation characterization of recombinant proteins.** SIDT1 and SIDT2 proteins recombinantly expressed in HEK293F cells with intact *N*-glycosylation was digested into peptides and the intact glycopeptides were analyzed with LC-MS/MS using EThcD mode.

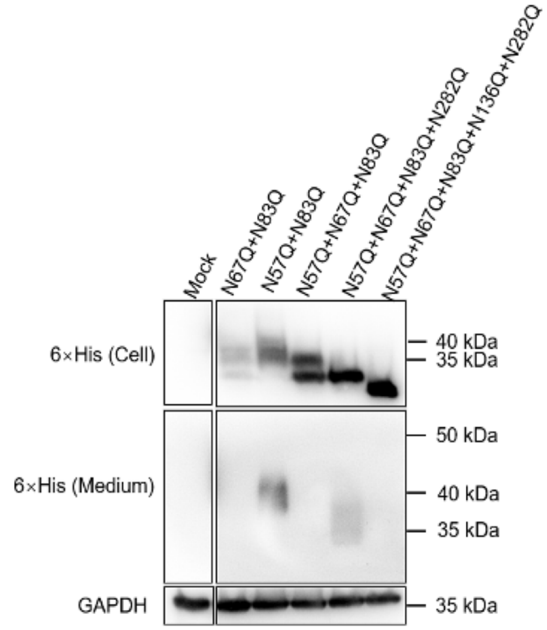


**Figure S3. Effect of *N*-glycosylation on SIDT1$^{ECD}$ protein secretion.** Western blots were analyzed for lysate and medium from SIDT1ECD and combined *N*-glycosites mutants transfected HEK293T cells. Three independent experiments were performed.

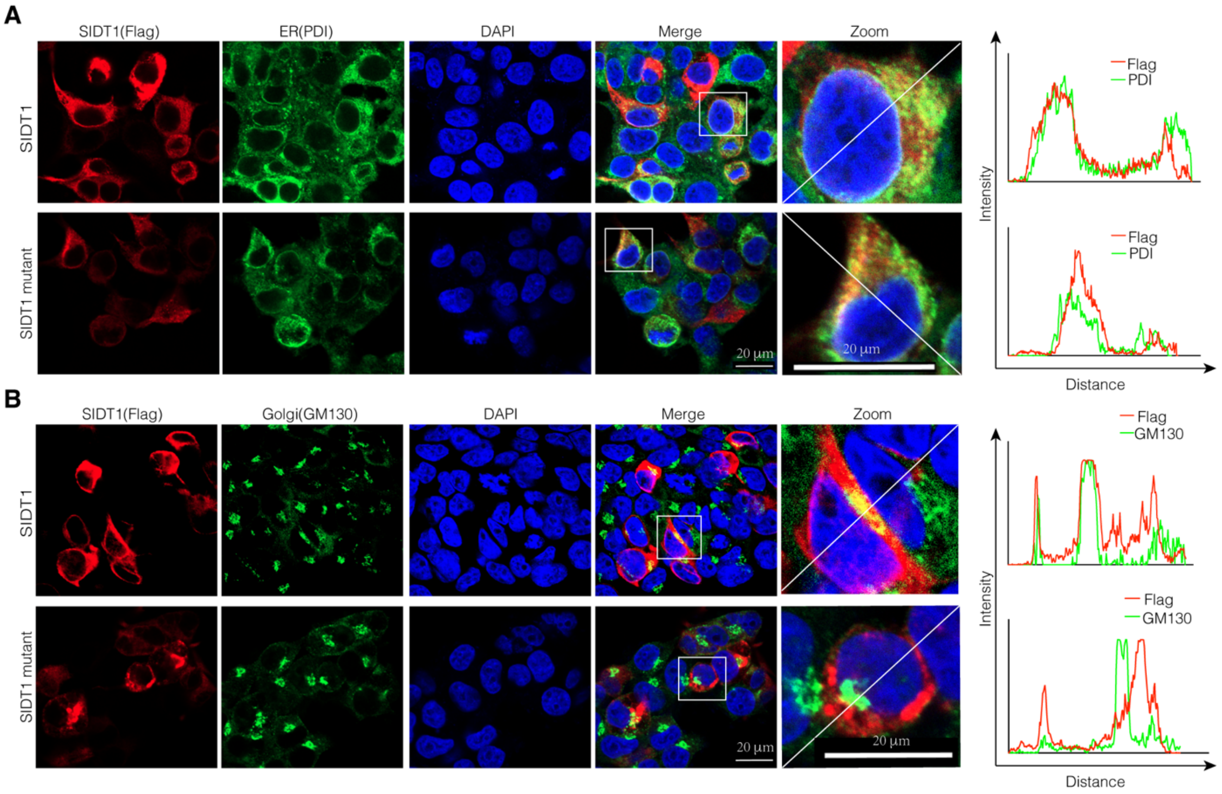


**Figure S4. The effect of *N*-glycosylation on the cell surface expression of SIDT1.** **A** Representative confocal microscopy images showing the ER localization in cells expressing wild-type SIDT1 and SIDT1 mutant variant. ER marker protein disulfide isomerase (PDI) (green) and anti-Flag (red) antibodies were used. Scale bar, 20 μm. **B** Representative confocal microscopy images showing the Golgi localization in cells expressing wild-type SIDT1 and SIDT1 mutant variant. Golgi marker GM130 (green) and anti-Flag (red) antibodies were used. Scale bar, 20 μm. Three independent experiments were performed.

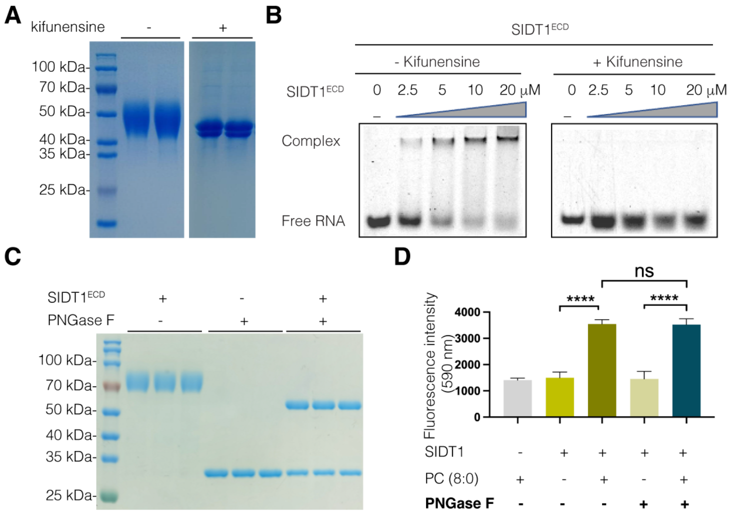


**Figure S5. Effect of *N*-glycosylation on RNA binding of SIDT1. A** SDS-PAGE analysis of purified kifunensine-treated and -untreated SIDT1$^{ECD}$. The Kifunensine-treated SIDT1$^{ECD}$ proteins were obtained by adding kifunensine during protein expression. **B** RNA binding of SIDT1$^{ECD}$ obtained from kifunensine-treated or -untreated HEK293F cells as determined by EMSA. The final protein concentrations of SIDT1$^{ECD}$ in lanes 1-5 are 0, 2.5, 5, 10, and 20 μM, respectively, and the final 5'-FAM-labeled ssRNA concentration is 2.5 μM. Complex: bound; Free ssRNA: unbound. **C** SDS-PAGE analysis of PNGase F-treated and -untreated SIDT1$^{ECD}$. Deglycosylated SIDT1$^{ECD}$ proteins were obtained by the deglycosylation with PNGase F. **D** Analysis of the phospholipase activity of PNGase F-treated and -untreated SIDT1. Deglycosylated SIDT1$^{ECD}$ proteins were obtained by the deglycosylation with PNGase F. Phospholipase activity was determined by comparing the fluorescence readings of test samples to a standard curve. Prism 8 software (GraphPad) was used to perform statistical analysis and determine significant differences between samples and controls. Three independent experiments were performed.

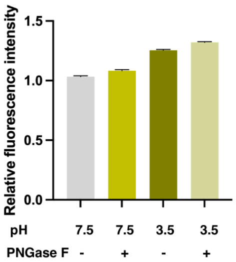


**Figure S6. Effect of *N*-glycosylation on SIDT1 mediated RNA uptake.** Cell viability of PNGase F-treated (+) and untreated (−) PANC1 cells was determined by CCK-8 assay. The optical density (OD) value of each well was measured at 450 nm. Three independent experiments were performed.
